# Mutation of influenza A virus PA-X decreases pathogenicity in chicken embryos and can increase the yield of reassortant candidate vaccine viruses

**DOI:** 10.1101/409375

**Authors:** Saira Hussain, Matthew L. Turnbull, Helen M. Wise, Brett W. Jagger, Philippa M. Beard, Kristina Kovacikova, Jeffery K. Taubenberger, Lonneke Vervelde, Othmar G Engelhardt, Paul Digard

**Author notes:** Address correspondence to Paul Digard. Present addresses: Saira Hussain, The Francis Crick Institute, London, United Kingdom. Matthew L Turnbull, Glasgow Centre for Virus Research, Glasgow, United Kingdom. Brett W. Jagger, Department of Medicine, Washington University in St. Louis, St. Louis, USA.

## Abstract

The PA-X protein of influenza A virus has roles in host cell shut-off and viral pathogenesis. While most strains are predicted to encode PA-X, strain-dependent variations in activity have been noted. We found that PA-X protein from A/PR/8/34 (PR8) strain had significantly lower repressive activity against cellular gene expression compared with PA-Xs from the avian strains A/turkey/England/50-92/91 (H5N1) (T/E) and A/chicken/Rostock/34 (H7N1). Loss of normal PA-X expression, either by mutation of the frameshift site or by truncating the X-ORF, had little effect on the infectious virus titre of PR8 or PR8 7:1 reassortants with T/E segment 3 grown in embryonated hens’ eggs. However, in both virus backgrounds, mutation of PA-X led to decreased embryo mortality and lower overall pathology; effects that were more pronounced in the PR8 strain than the T/E reassortant, despite the low shut-off activity of the PR8 PA-X. Purified PA-X mutant virus particles displayed an increased ratio of HA to NP and M1 compared to their WT counterparts, suggesting altered virion composition. When the PA-X gene was mutated in the background of poorly growing PR8 6:2 vaccine reassortant analogues containing the HA and NA segments from H1N1 2009 pandemic viruses or an avian H7N3 strain, HA yield increased up to 2-fold. This suggests that the PR8 PA-X protein may harbour a function unrelated to host cell shut-off and that disruption of the PA-X gene has the potential to improve the HA yield of vaccine viruses.

**IMPORTANCE:** Influenza A virus is a widespread pathogen that affects both man and a variety of animal species, causing regular epidemics and sporadic pandemics with major public health and economic consequences. A better understanding of virus biology is therefore important. The primary control measure is vaccination, which for humans, mostly relies on antigens produced in eggs from PR8-based viruses bearing the glycoprotein genes of interest. However, not all reassortants replicate well enough to supply sufficient virus antigen for demand. The significance of our research lies in identifying that mutation of the PA-X gene in the PR8 strain of virus can improve antigen yield, potentially by decreasing the pathogenicity of the virus in embryonated eggs.

## Introduction

Influenza epidemics occur most years as the viruses undergo antigenic drift. Influenza A viruses (IAV) and influenza B viruses cause seasonal human influenza but IAV poses an additional risk of zoonotic infection, with the potential of a host switch and the generation of pandemic influenza. The 1918 ‘Spanish flu’ pandemic was by far the worst, resulting in 40-100 million deaths worldwide (1), while the 2009 swine flu pandemic caused an estimated 200,000 deaths worldwide (2).

IAV contains eight genomic segments encoding for at least ten proteins. Six genomic segments (segments 1, 2, 3, 5, 7 and 8) encode the eight core “internal” proteins PB2, PB1, PA, NP, M1, NS1 and NS2, as well as the ion channel M2. These segments can also encode a variety of accessory proteins known to influence pathogenesis and virulence (reviewed in (3, 4)). Segments 4 and 6 encode for the two surface glycoproteins haemagglutinin (HA) and neuraminidase (NA) respectively (5, 6) and virus strains are divided into subtypes according to the antigenicity of these proteins.

Vaccination is the primary public health measure to reduce the impact of influenza epidemics and pandemics, principally using inactivated viruses chosen to antigenically match the currently circulating virus strains or newly emerging viruses of pandemic concern. However, before efficient vaccine production can commence, high-yielding candidate vaccine viruses (CVVs) need to be prepared. Seasonal CVVs are widely produced by classical reassortment. This process involves co-infecting embryonated hens’ eggs with the vaccine virus along with a high yielding “donor” virus adapted to growth in eggs (most commonly the A/Puerto Rico/8/34 strain, or “PR8”). The highest yielding viruses that contain the glycoproteins of the vaccine virus are then selected. Recombinant influenza viruses are also made by reverse genetics (RG) (7-9), which relies on the transfection of cells with plasmids engineered to express both viral genomic RNA and proteins from each of the eight segments and hence initiate virus production; the resultant virus is subsequently amplified in eggs. When making RG CVVs, typically the six segments encoding core proteins (backbone) are derived from the donor strain whereas the two segments encoding the antigens are derived from the vaccine virus. Classical reassortment has the advantage that it allows for the fittest natural variant to be selected but it can be time consuming. In the case of a pandemic, large quantities of vaccine must be made available quickly. Moreover, RG is the only viable method for production of CVVs for potentially pandemic highly pathogenic avian influenza viruses, since it allows for removal of genetic determinants of high pathogenicity in the virus genome, as vaccines are manufactured in biosafety level 2 laboratories. A limited number of donor strains for IAV vaccine manufacture currently exist. Although PR8 is widely used, reassortant viruses based on it do not always grow sufficiently well for efficient vaccine manufacture. In the case of the 2009 H1N1 pandemic (pdm09), vaccine viruses grew poorly in eggs compared with those for previous seasonal H1N1 isolates (10), resulting in manufacturers struggling to meet demand. Thus, there is a clear need for new reagents and methods for IAV production, particularly for pandemic response.

In recent years, several approaches have been employed to improve antigen yield of candidate vaccine viruses made by reverse genetics. These have involved empirical testing and selection of PR8 variants (11, 12), as well as targeted approaches such as making chimeric genes containing promoter and packaging signal regions of PR8 while encoding the ectodomain of the CVV glycoprotein genes (13-21), or introducing a wild-type (WT) virus-derived segment 2 (21-29). Our approach was to manipulate expression of an accessory protein virulence factor, PA-X (30). Segment 3, encoding PA as the primary gene product, also expresses PA-X by low-level ribosomal shifting into a +1 open reading frame (ORF) termed the X ORF (Fig 1A) (30). PA-X is a 29 kDa protein that contains the N-terminal endonuclease domain of PA, and in most isolates, a 61 amino acid C-terminus from the X ORF (30-32). It has roles in shutting off host cell protein synthesis and, at the whole animal level, modulating the immune response (30, 33). Loss of PA-X expression has been shown to be associated with increased virulence in mice for 1918 H1N1, H5N1 and also pdm09 and classical swine influenza H1N1 strains, as well as in chickens and ducks infected with a highly pathogenic H5N1 virus (30, 34-40). However, in other circumstances, such as avian H9N2 viruses (40) or, in some cases, A(H1N1)pdm09 viruses (37, 41), mutation of PA-X resulted in reduced pathogenicity in mice. Similarly, a swine influenza H1N2 virus (42) lacking PA-X showed reduced pathogenicity in pigs. Moreover, PA-X activity in repressing cellular gene expression is strain dependent (33, 34, 40, 43), with laboratory-adapted viruses such as A/WSN/33 showing lower levels of activity (33). Here, we show that although the PR8 PA-X polypeptide has low shut-off activity, removing its expression decreases the pathogenicity of the virus in the chick embryo model. Moreover, we found that, for certain poor growing CVV mimics, ablating PA-X expression improved HA yield from embryonated eggs up to 2-fold. In no case did loss of PA-X appear to be detrimental to the growth of CVVs, making it a potential candidate mutation for incorporation into the PR8 CVV donor backbone.

**FIGURE 1.**
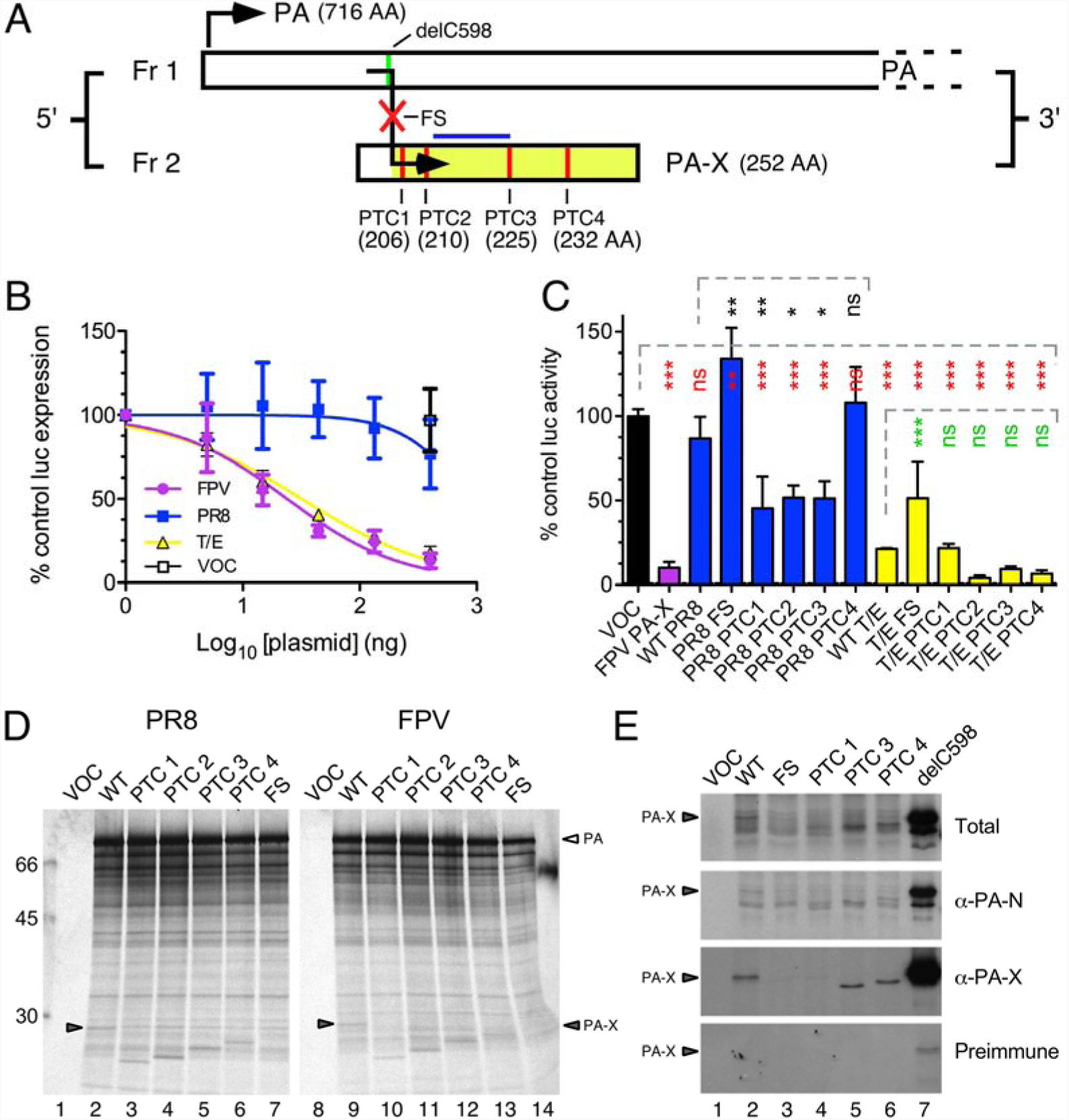
Virus strain dependent variation in PA-X-mediated host cell shut-off activity. **A)** Schematic showing mutations in segment 3: at the frameshift (FS) site to generate a PA-X null virus, or in the X-ORF so that segment 3 expresses C-terminally truncated versions of PA-X (PTCs 1-4, size of products indicated), or removing cytosine 598 (delC598) to place the X ORF in frame with PA such that only PA-X is expressed. **B, C)** PA-X-mediated inhibition of cellular RNA polymerase II-driven gene expression in QT-35 cells. **B)** Cells were co-transfected with 100 ng of pRL plasmid constitutively expressing *Renilla* luciferase and a dilution series of the indicated segment 3 pHW2000 plasmids or with a fixed amount of the empty pHW2000 vector (VOC). Luciferase activity was measured 48 h later and plotted as % of a pRL only sample. Dose-inhibition curves were fitted using GraphPad Prism software. Data are mean ± SD of two independent experiments each performed in triplicate. **C)** Cells were co-transfected with 100 ng of pRL plasmid and 400 ng of effector pHW2000 plasmids expressing segment 3 products. Luciferase activity was measured 48 h later and plotted as the % of a pHW2000 vector only control. Data are the mean ± SD from 2 independent experiments performed in duplicate. Dashed lines indicate groups of statistical tests (against the left hand bar in each case; * p < 0.05, ** p < 0.01, *** p < 0.001) as assessed by Dunnett’s test. **D, E)** *In vitro* translation of PA-X from PR8 segment 3 constructs. Aliquots of rabbit reticulocyte lysate supplemented with ^35^S-methionine were programmed with the indicated plasmids and radiolabelled polypeptides visualised by SDS-PAGE and autoradiography before (**D**) or after (**E**) immunoprecipitation with the indicated antisera. Arrowheads in (D) indicate full length PA-X while molecular mass (kDa) markers are shown on the left.

## Results

### The PR8 virus strain PA-X has relatively low shut-off activity

Previous work has noted variation in apparent activity of PA-X proteins from different strains of virus, with the laboratory adapted-strain WSN showing lower activity than many other strains (33). Re-examination of evidence concerning a postulated proteolytic activity of PA (43) suggested that lower PA-X activity might also be a feature of the PR8 strain. To test this, the ability of PR8 segment 3 gene products to inhibit cellular gene expression was compared to that of two avian virus-derived PA segments (from A/chicken/Rostock/34 [H7N1; FPV] and A/turkey/England/50-92/91 [H5N1; T/E]. Avian QT-35 (Japanese quail fibrosarcoma) cells were transfected with a consistent amount of a plasmid encoding luciferase under the control of a constitutive RNA polymerase II promoter (pRL) and increasing amounts of the IAV cDNAs (in pHW2000-based RG plasmids), or as a negative control, the maximum amount of the empty pHW2000 vector. Luciferase expression was measured 48 h later and expressed as a % of the amount obtained from pRL-only transfections. Transfection of a 4-fold excess of empty pHW2000 vector over the luciferase reporter plasmid had no significant effect on luciferase expression, whereas co-transfection of the same amount of either the FPV or T/E segments suppressed activity to around 10% of the control (Fig 1B). Titration of the FPV and T/E plasmids gave a clear dose-response relationship, giving estimated EC_50_ values of 24 ± 1.1 ng and 32 ± 1.1 ng respectively. In contrast, the maximum amount of the PR8 plasmid only inhibited luciferase expression by around 30% and an EC_50_ value could not be calculated, indicating a lower ability to repress cellular gene expression. Similarly low inhibitory activity of the PR8 segment 3 was seen in a variety of other mammalian cell lines (data not shown), suggesting it was an intrinsic feature of the viral gene, rather than a host- or cell type-specific outcome.

Several studies have shown the X-ORF to be important in host cell shut-off function and virulence of PA-X (37, 44-46). To further explore the influence of X-ORF sequences on virus strain-specific host cell shut-off, mutations were constructed in segment 3 in which PA-X expression was either hindered (*via* mutation of the frameshift site [FS]) or altered by the insertion of premature termination codons (PTCs 1-4; silent in the PA ORF) such that C-terminally truncated forms of PA-X would be expressed (Figure 1A). QT-35 cells were co-transfected with the pRL plasmid and a fixed amount of WT, FS or PTC plasmids and luciferase expression measured 48 h later. As before, the WT FPV and T/E PA-Xs reduced luciferase activity by approximately 5-10 fold, while WT PR8 PA-X had no significant effect (Figure 1C). Introducing the FS mutation into both PR8 and T/E segment 3 significantly increased luciferase activity relative to the WT construct. Truncation of the PR8 PA-X to 225 AA or less (PTC mutations 1-3) significantly improved shut-off activity, although not to the levels seen with the WT avian virus polypeptides, while the PTC4 truncation had no effect. In contrast, none of the PTC mutations significantly affected activity of the T/E PA-X, although there was a trend towards increased activity from the PTC2, 3 and 4 truncations

Low activity could be due to decreased expression and/or decreased activity of PA-X. To examine this, expression of the low activity PR8 and high activity FPV PA-X constructs were compared by *in vitro* translation reactions in rabbit reticulocyte lysate. Translation of segment 3 from both PR8 and FPV produced both full length PA and similar quantities of a minor polypeptide species of the expected size for PA-X whose abundance decreased after addition of the FS mutation or whose electrophoretic mobility was altered in stepwise fashion after C-terminal truncation with the PTC1-4 mutations (Figure 1D). This suggested that differences in shut-off potential were not linked to intrinsic differences in PA-X protein synthesis. To confirm the identity of the PR8 *in vitro* translated polypeptides, immunoprecipitation of IVT products with sera raised either against the N-terminal domain of PA, or an X-ORF derived polypeptide or pre-immune sera (30) were performed (Figure 1E). WT PA-X was clearly visible in samples immunoprecipitated with anti-PA-X and anti-PA-N but not the pre-immune serum, where it co-migrated with the product from the delC598 plasmid, a construct in which cytosine 598 of segment 3 (the nucleotide skipped during the PA-X frameshifting event (47)) had been deleted to put the X-ORF into the same reading frame as the N-terminal PA domain (Figure 1E, lanes 2 and 7). In contrast, only background amounts of protein were precipitated from the FS IVT (lane 3). Faster migrating polypeptide products from the PTC3 and 4 plasmids showed similar reactivities to WT PA-X (lanes 5 and 6) whereas the product of the PTC1 plasmid was only precipitated by anti-PA-N (lane 4), as expected because of the loss of the epitope used to raise the PA-X antiserum (Figure 1A). Overall therefore, the PR8 PA-X polypeptide possessed lower shut-off activity than two avian virus PA-X polypeptides despite comparable expression *in vitro*, and its activity could be modulated by mutation of the X-ORF.

### Loss of PA-X expression results in significantly less pathogenicity in chick embryos without affecting virus replication

In order to further characterise the role of PA-X as a virulence determinant, we tested the panel of high and low activity mutants in the chicken embryo pathogenicity model. Embryonated hens’ eggs were infected with PR8-based viruses containing either PR8 or T/E WT or mutant segment 3s and embryo viability was monitored at 2 days post infection (p.i.) by candling. Both WT PR8 and the WT 7:1 reassortant with the T/E segment 3 viruses had killed over 50% of the embryos by this point (Figures 2A and B). Truncation of PA-X by the PTC mutations led to small improvements in embryo survival, although none of the differences were statistically significant. However, embryo lethality was significantly reduced to below 20% following infection with the PR8 FS virus compared with PR8 WT virus. A similar reduction in lethality was seen for the T/E FS virus, although the difference was not statistically significant. This reduction in embryo pathogenicity following ablation of PA-X expression suggested potential utility as a targeted mutation in the PR8 backbone used to make CVVs. Accordingly, to characterise the effects of mutating PR8 PA-X over the period used for vaccine manufacture, embryo survival was monitored daily for 72 h. Eggs infected with WT PR8 showed 45% embryo survival at 2 days p.i. and all were dead by day 3 (Figure 2C). However, the PR8 FS infected eggs showed a statistically significant improvement in survival compared to WT, with 80% and 30% survival at days 2 and 3, respectively. Embryos infected with PR8 expressing the C-terminally truncated PTC1 form of PA-X showed an intermediate survival phenotype with 60% and 20% survival at days 2 and 3, respectively.

**FIGURE 2.**
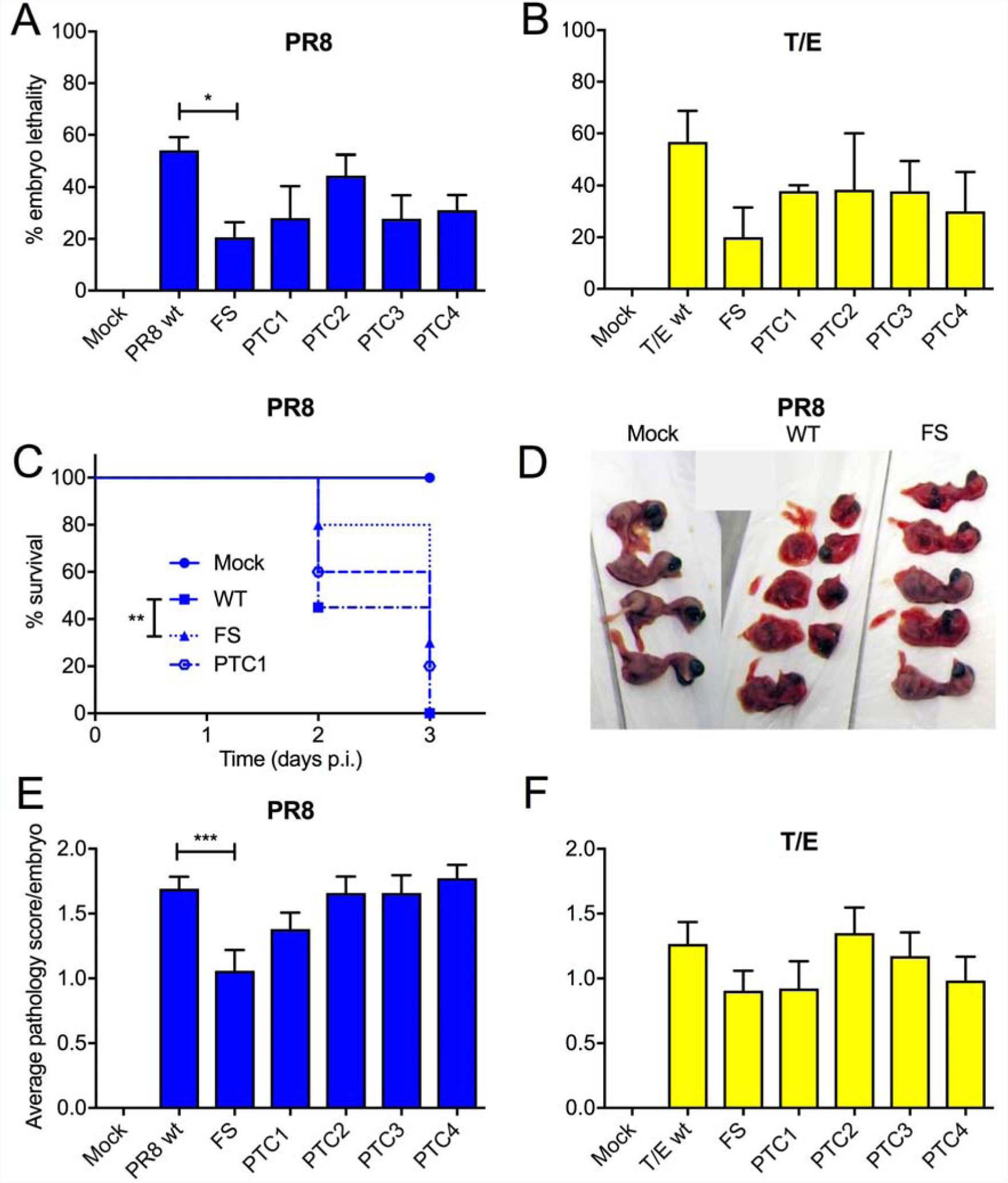
Effect of PA-X mutations in a chick embryo pathogenicity model. Groups of 5-6 embryonated hens’ eggs were infected with 1000 PFU of the indicated viruses and **A, B**) embryo viability was determined by candling at 2 days p.i. Data are plotted as mean ± SEM % embryo lethality from 3-4 independent experiments. Horizontal bars indicate statistical significance (* p < 0.05) as assessed by Dunnett’s test. **C)** Infected eggs were monitored daily for embryo viability and survival was plotted versus time. Data are from 3 independent experiments with 5 - 10 eggs per experiment. Statistical significance between WT and FS viruses (** p < 0.01) was assessed by log-rank (Mantel-Cox) test. **D-F)** From the experiments described in **A)** and **B)**, embryos were imaged **D)** and **E, F)** scored blind by two observers as 0 = normal, 1= intact but bloody, 2 = small, damaged and with severe haemorrhages. Data are the average ± SEM pathology scores from 3-4 independent experiments. Horizontal bar indicates statistical significance (*** p < 0.001) as assessed by Dunnett’s test.

To further assess the effects of mutating PA-X, the chicken embryos were examined for gross pathology. WT PR8 infection resulted in smaller, more fragile embryos with diffuse reddening, interpreted as haemorrhages (Figure 2D). In comparison, the PA-X null FS mutant-infected embryos remained intact, were visibly larger and less red. To quantitate these observations, embryos were scored blind for gross pathology. Taking uninfected embryos as a baseline, it was clear that WT PR8 virus as well as the PA-X truncation mutants induced severe changes to the embryos (Figure 2E). In contrast, the PA-X null FS mutant caused significantly less pathology. The WT 7:1 T/E reassortant virus gave less overt pathology than WT PR8 but again, reducing PA-X expression through the FS mutation further reduced damage to the embryos (Figure 2E). Similar trends in pathology were also seen with 7:1 PR8 reassortant viruses containing either WT or FS mutant versions of FPV segment 3 (data not shown).

Examination of haematoxylin and eosin (H&E) stained sections through the embryos revealed pathology in numerous organs including the brain, liver and kidney (Figure 3). In the brain of embryos infected with WT virus there was marked rarefaction of the neuropil, few neurons were identifiable, and there was accumulation of red blood cells (Figure 3C). In the liver of embryos infected with WT virus the hepatic cords were disorganised, and the hepatocytes were often separated by large pools of red blood cells (Figure 3D). In the kidney of embryos infected with WT virus, tubules were often lined by degenerate epithelial cells (characterised by loss of cellular detail). In all cases the pathology noted in WT virus-infected embryos was also present in the FS virus-infected embryos but at a reduced severity. Thus overall, disruption of PA-X expression in PR8 resulted in significantly less pathogenicity in chick embryos.

**FIGURE 3.**
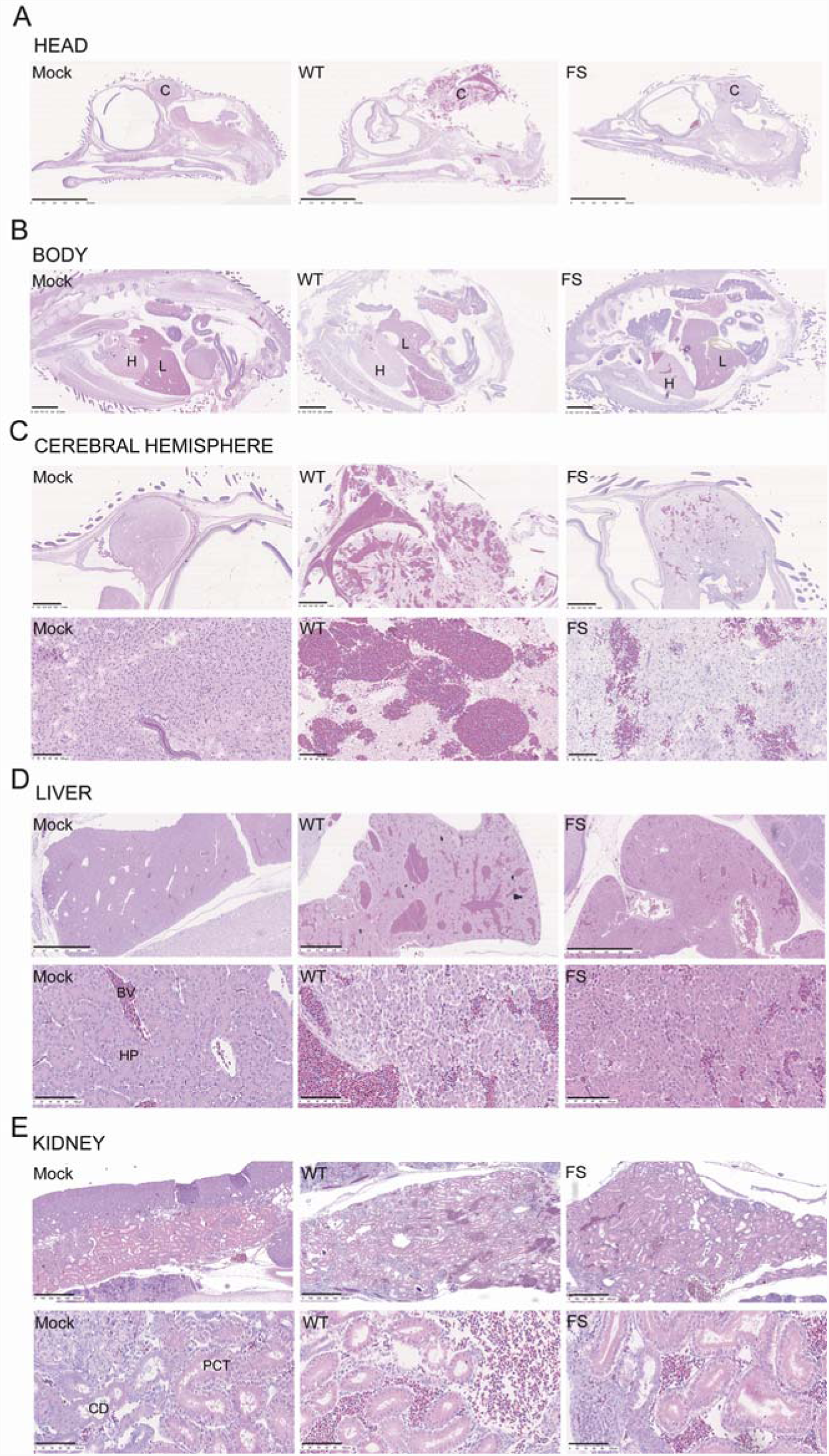
Histopathology of chick embryos following infection with PR8 viruses. Embryonated hens’ eggs were infected with segment 3 WT or mutant viruses or mock infected. 2 days p.i. embryos were fixed, sectioned and stained with H&E before imaging with a Nanozoomer XR (Hamamatsu) using brightfield settings; representative pictures are shown: **A)** head, scale bar = 5 mm, and **B)** body, scale bar = 2.5 mm, and **C)** cerebral hemisphere, **D)** liver and **E)** kidney, scale bars for low and high magnification images = 1mm and 100 μm or 500 μm, respectively. C = cerebral hemisphere, H = heart, L = liver, PCT = proximal convoluted tubule, CD = collecting duct, HP=hepatocytes, BV= blood vessel).

Reduced pathogenicity *in vivo* following loss of PA-X expression could be due to a replication deficiency of the virus, although the viruses replicated equivalently in mammalian MDCK cells (data not shown). To test if replication did differ *in ovo*, infectious virus titres were obtained (by plaque titration on MDCK cells) from the allantoic fluid of embryonated hens’ eggs infected with the panels of PR8 and T/E viruses at 2 days p.i.. However, there were no significant differences in titres between either PR8 or T/E WT and PA-X mutant viruses (Figures 4A, B). Since the reduced pathogenicity phenotype *in ovo* on loss of PA-X expression was more pronounced for viruses with PR8 segment 3 than the T/E gene, embryos from PR8 WT and segment 3 mutant-infected eggs were harvested at 2 days p.i., washed, macerated and virus titres from the homogenates determined. Titres from embryos infected with the PR8 FS and PTC4 viruses were slightly (less than 2-fold) reduced compared to embryos infected with PR8 WT virus (Figure 4C), but overall there were no significant differences in titres between the viruses. To see if there were differences in virus localisation in tissues between PR8 WT and FS viruses, immunohistochemistry was performed on chick embryo sections to detect viral NP as a marker of infected cells. NP positive cells were seen in blood vessels throughout the head and body of both PR8 WT and FS-infected embryos; liver, heart and kidney are shown as representatives (Figure 4D), indicating that the circulatory system had been infected. However, there were no clear differences in virus localisation between embryos infected with WT and FS viruses.

**FIGURE 4.**
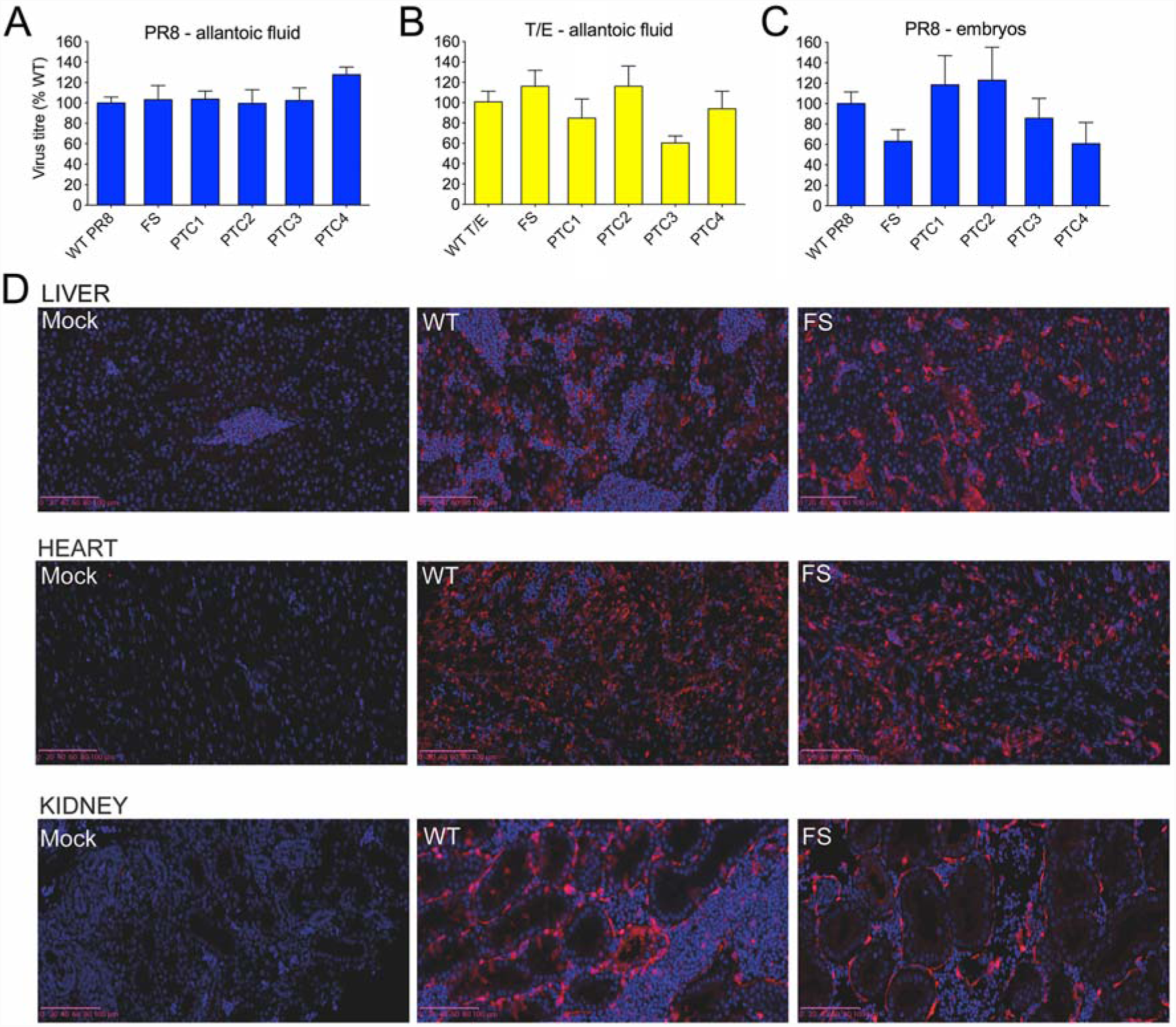
Effects of mutating PA-X expression on virus replication in chick embryos. Groups of 5-6 embryonated hens’ eggs were infected with the indicated viruses and at 2 days p.i., virus titres determined by plaque assay from (**A, B**) allantoic fluid or (**C)** washed and macerated chick embryos. Graphs represent the mean ± SEM from 3 (A, B) or 2-4 independent experiments (C). Titres of mutant viruses were not significantly different compared to WT virus (Dunnett’s test). **D)** Embryos were fixed at 2 days p.i., sectioned and stained for IAV NP and DNA before imaging using a Nanozoomer XR (Hamamatsu) on fluorescence settings. Representative images of liver, heart and kidney are shown. Scale bars = 100 μm. NP = red, DAPI = blue.

Overall therefore, the loss of PA-X expression reduced IAV pathogenicity in chick embryos, as assessed by mortality curves and both gross and histopathological examination of embryo bodies. This reduced pathogenicity did not appear to correlate with reduced replication or altered distribution of the virus *in ovo*.

### Ablating PA-X expression alters virion composition

Other viruses encode host-control proteins with mRNA endonuclease activity, including the SOX protein of murine gammaherpesvirus MHV68 whose expression has been shown to also modulate virion composition (48). Also, egg-grown IAV titre and HA yield do not always exactly match, with certain problematic candidate vaccine viruses (CVVs) containing lower amounts of HA per virion (16, 49, 50). Accordingly, we compared the relative quantities of virion structural proteins between PA-X expressing and PA-X null viruses. Two pairs of viruses were tested: either an entirely PR8-based virus, or a 7:1 reassortant of PR8 with FPV segment 3, both with or without the FS mutation. Viruses were grown in eggs as before and purified from allantoic fluid by density gradient ultracentrifugation before polypeptides were separated by SDS-PAGE and visualised by staining with Coomassie blue. To ensure that overall differences in protein loading did not bias the results, 3-fold dilutions of the samples were analysed. From the gels, the major virion components of both WT and FS virus preparations could be distinguished: NP, the two cleaved forms of haemagglutinin, HA1 and HA2, the matrix protein, M1 and in lower abundance, the polymerase proteins (Figures 5A, B, lanes 4-9). In contrast, only trace polypeptides were present in similarly purified samples from uninfected allantoic fluid (lanes 1-3). Densitometry was used to assess the relative viral protein contents of the viruses. The two most heavily loaded lanes (where band intensities were sufficient for accurate measurement) were quantified and average HA1:NP and HA2:M1 ratios calculated. When the data from three independent experiments were examined in aggregate by scatter plot (Figures 5C and D), a statistically significant increase in the average quantity of HA1 relative to NP was evident for both PR8 and the FPV reassortant FS viruses of ∼1.4-fold and ∼1.6-fold respectively compared to WT (Figures 5 C, D). The ratio of HA2:M1 was also significantly increased in the PR8 FS virus (∼1.2-fold greater for WT) and a similar but non-significant increase was seen for the FPV virus pair. These data are consistent with the hypothesis that PA-X expression modulates virion composition.

**FIGURE 5.**
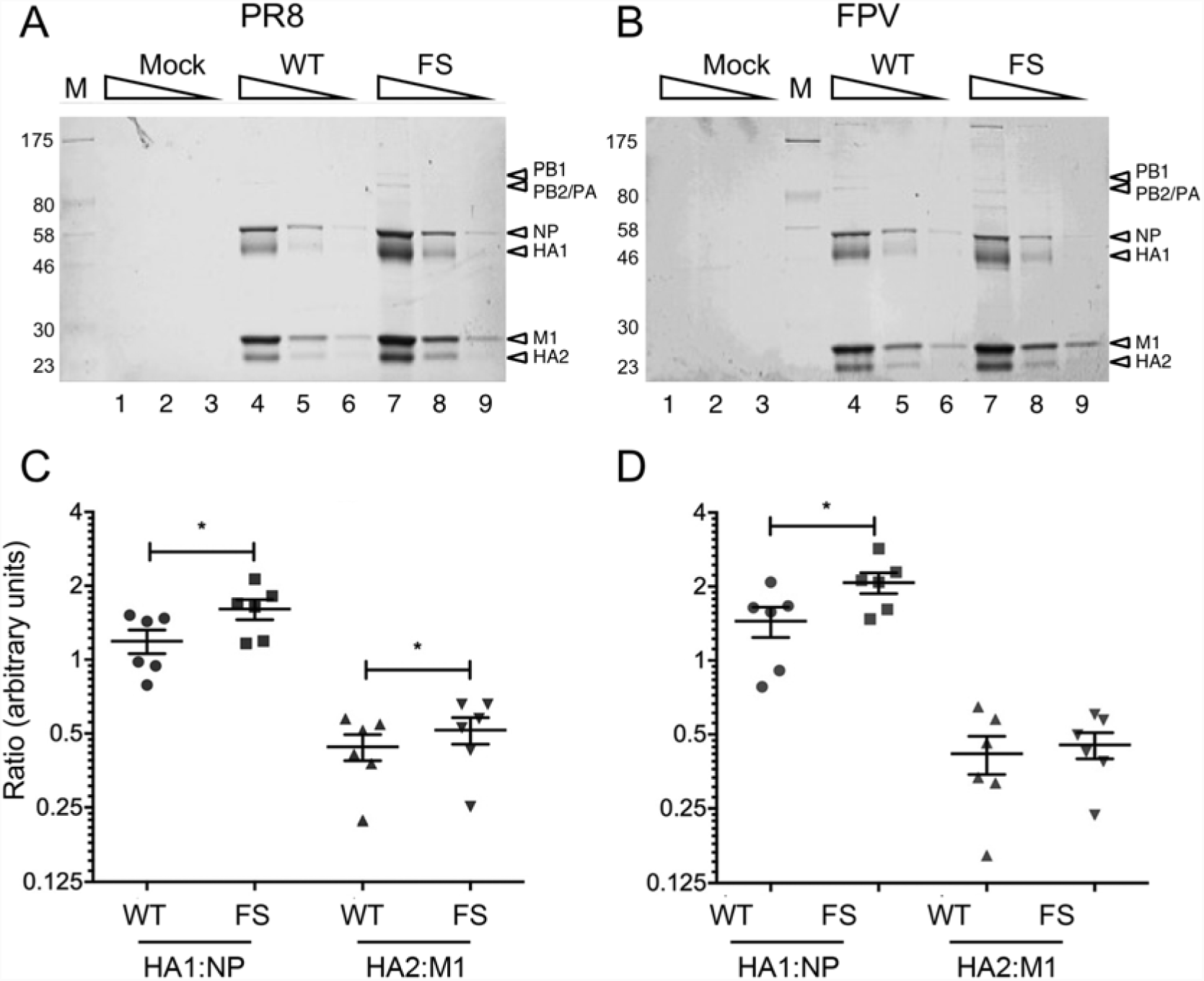
Virion composition of WT or FS mutant viruses. Embryonated hens’ eggs were infected with WT or segment 3 mutant viruses or mock infected. At 2 days p.i., virus was purified from allantoic fluid by sucrose density gradient ultracentrifugation and 3-fold serial dilutions **A, B)** analysed by SDS-PAGE on 10% polyacrylamide gels and staining with Coomassie blue. **C, D)** For PR8 and FPV, respectively, the ratios of NP:HA1 and M1:HA2 were determined by densitometry of SDS-PAGE gels. Scatter plots with the mean and SEM of 6 measurements from 3 independent experiments using 2 independently rescued virus stocks are shown. Horizontal bars indicate statistical significance (* p < 0.05) as assessed by paired t-test.

### Ablating PA-X expression increases HA yield of CVVs bearing pdm2009 glycoproteins

The reduced pathogenicity and corresponding longer embryo survival time induced by the PR8 FS mutant *in ovo* coupled with evident modulation of virion composition in favour of HA content suggested a strategy to increase overall antigen yields for PR8-based CVVs. Therefore, the effect of incorporating the PA-X FS mutation into CVV mimics containing glycoproteins of different IAV subtypes was examined. Reasoning that a benefit might be most apparent for a poor-yielding strain, 6:2 CVV mimics containing the glycoprotein genes from the A(H1N1)pdm09 vaccine strain, A/California/07/2009 (Cal7) with the six internal genes from PR8, with or without the FS mutation in segment 3, were generated. Growth of these viruses in embryonated hens’ eggs was then assessed by inoculating eggs with either 100, 1,000 or 10,000 PFU per egg (modelling the empirical approach used in vaccine manufacture to find the optimal inoculation dose) and measuring HA titre at 3 days p.i.. Both viruses grew best at an inoculation dose of 100 PFU/egg, but final yield was both relatively low (as expected, ∼ 64 HAU/50 μl) and insensitive to input dose, with average titres varying less than 2-fold across the 100-fold range of inocula (Figure 6A). However, at each dose, the 6:2 FS virus gave a higher titre (on average, 1.6-fold) than the parental 6:2 reassortant. In order to assess HA yield between the WT and FS viruses on a larger scale, comparable to that used by WHO Essential Regulatory Laboratories (ERLs) such as the National Institute for Biological Standards and Control, UK, 20 eggs per virus were infected at a single inoculation dose. In this experiment, the average HA titre of the FS virus was over 3 times higher than the WT 6:2 virus (Figure 6B). To further determine the consistency of these results, HA titre yields were assessed from two independently rescued reverse genetics stocks of the Cal7 6:2 CVV mimics with or without the PR8 PA-X gene as well as another 6:2 CVV mimic bearing the glycoproteins from the A/England/195/2009 (Eng195) A(H1N1)pdm09 strain. HA yield from different stocks was assessed in independent repeats of both small-(5 eggs for each of three different inoculation doses, taking data from the dose that gave maximum yield) and large-scale (20 eggs per single dose of virus) experiments. Examination of the average HA titres showed considerable variation between independent experiments (Figure 6C). However, when plotted as paired data points, it was obvious that in every experiment, the FS viruses gave a higher yield than the parental 6:2 reassortant and on average, there were 2.7-and 3.8-fold higher HA titres with the segment 3 FS mutation for Cal7 and Eng195 respectively (Table 1).

**TABLE 1.**
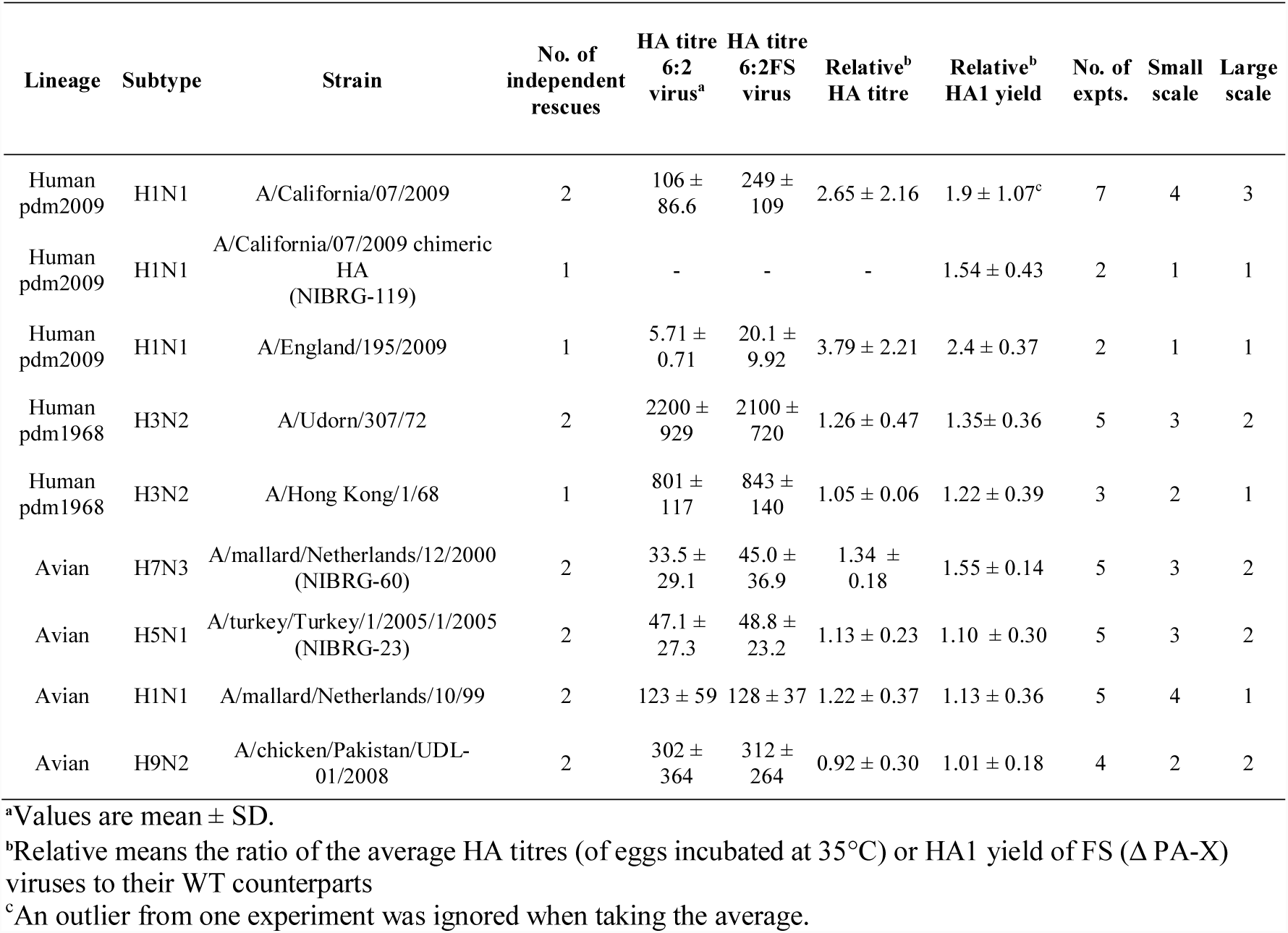
**Effects of the** Δ**PA-X FS mutation on HA yield of CVVs grown in eggs.**

| Lineage | Subtype | Strain | No. of independent rescues | HA titre 6:2 virus<sup>a</sup> | HA titre 6:2FS virus | Relative<sup>b</sup> HA titre | Relative<sup>b</sup> HA1 yield | No. of expts. | Small scale | Large scale |
| --- | --- | --- | --- | --- | --- | --- | --- | --- | --- | --- |
| Human pdm2009 | H1N1 | A/California/07/2009 | 2 | 106 ± 86.6 | 249 ± 109 | 2.65 ± 2.16 | 1.9 ± 1.07<sup>c</sup> | 7 | 4 | 3 |
| Human pdm2009 | H1N1 | A/California/07/2009 chimeric HA (NIBRG-119) | 1 | - | - | - | 1.54 ± 0.43 | 2 | 1 | 1 |
| Human pdm2009 | H1N1 | A/England/195/2009 | 1 | 5.71 ± 0.71 | 20.1 ± 9.92 | 3.79 ± 2.21 | 2.4 ± 0.37 | 2 | 1 | 1 |
| Human pdm1968 | H3N2 | A/Udorn/307/72 | 2 | 2200 ± 929 | 2100 ± 720 | 1.26 ± 0.47 | 1.35 ± 0.36 | 5 | 3 | 2 |
| Human pdm1968 | H3N2 | A/Hong Kong/1/68 | 1 | 801 ± 117 | 843 ± 140 | 1.05 ± 0.06 | 1.22 ± 0.39 | 3 | 2 | 1 |
| Avian | H7N3 | A/mallard/Netherlands/12/2000 (NIBRG-60) | 2 | 33.5 ± 29.1 | 45.0 ± 36.9 | 1.34 ± 0.18 | 1.55 ± 0.14 | 5 | 3 | 2 |
| Avian | H5N1 | A/turkey/Turkey/1/2005/1/2005 (NIBRG-23) | 2 | 47.1 ± 27.3 | 48.8 ± 23.2 | 1.13 ± 0.23 | 1.10 ± 0.30 | 5 | 3 | 2 |
| Avian | H1N1 | A/mallard/Netherlands/10/99 | 2 | 123 ± 59 | 128 ± 37 | 1.22 ± 0.37 | 1.13 ± 0.36 | 5 | 4 | 1 |
| Avian | H9N2 | A/chicken/Pakistan/UDL-01/2008 | 2 | 302 ± 364 | 312 ± 264 | 0.92 ± 0.30 | 1.01 ± 0.18 | 4 | 2 | 2 |

<sup>a</sup>Values are mean ± SD.
<sup>b</sup>Relative means the ratio of the average HA titres (of eggs incubated at 35°C) or HA1 yield of FS ( ΔPA-X) viruses to their WT counterparts
<sup>c</sup>An outlier from one experiment was ignored when taking the average.

**FIGURE 6.**
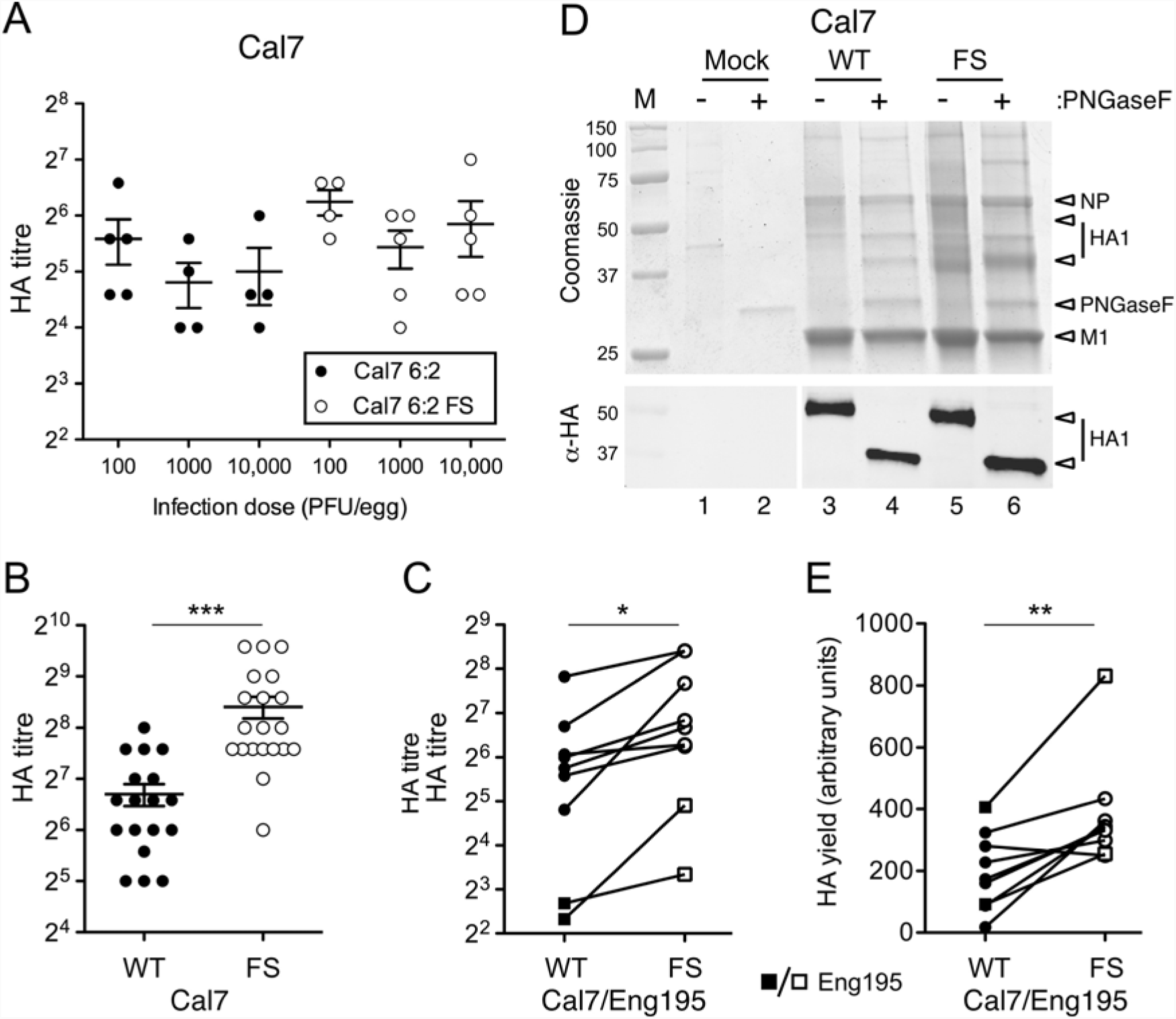
Effect of the PA-X FS mutation on HA yield of A(H1N1) pdm09 CVV mimics. Embryonated hens’ eggs were infected as indicated and **A-C)** HA titres in allantoic fluid measured at 3 days post-infection for (A) groups of 5 or (B) 20 eggs per condition. (**A, B**) Scatter plots of titres from individual eggs with mean and SEM are shown. (*** p < 0.001) assessed by unpaired t-test. **C)** Average HA titres from groups of eggs inoculated at the infection dose which gave maximum yield are shown as paired observations. Statistical significance (* p < 0.05, n=9) assessed by paired t-test. **D, E)** Allantoic fluid was clarified and virus pelleted by ultracentrifugation through 30% sucrose pads. Equal volumes of resuspended virus pellets were separated by SDS-PAGE on a 12% polyacrylamide gel and visualized by (**D**) staining with Coomassie blue (upper panel) or western blot for HA1 (lower panel) with (+) or without (-) prior treatment with PNGase F. Molecular mass (kDa) markers and specific polypeptides are labelled. **E)** De-glycosylated HA1 yield was quantified by densitometry of the western blots. Data points represent 8 independent experiments using 3 independently rescued RG virus stocks shown as paired observations. (** p < 0.01, n=8) as assessed by paired t-test. Circles represent Cal7 and squares represent Eng195 CVV mimics.

To directly assess HA protein yield, viruses were partially purified by ultracentrifugation of pooled allantoic fluid through 30% sucrose cushions. Protein content was analysed by SDS-PAGE and Coomassie staining, either before or after treatment with N-glycosidase F (PNGaseF) to remove glycosylation from HA and NA. Both virus preparations gave polypeptide profiles that were clearly different from uninfected allantoic fluid processed in parallel, with obvious NP and M1 staining, as well other polypeptide species of less certain origin (Figure 6D). Overall protein recovery was higher in the FS virus than the WT reassortant virus (compare lanes 3 and 4 with 5 and 6), but the poor yields of these viruses made unambiguous identification of the HA polypeptide difficult. However, PNGaseF treatment led to the appearance of a defined protein band migrating at around 40 kDa that probably represented de-glycosylated HA1, and this was present in appreciably higher quantities in the 6:2 FS preparation (compare lanes 4 and 6). Therefore, equivalent amounts of glycosylated or de-glycosylated samples from the Cal7 WT and FS reassortants were analysed by SDS-PAGE and western blotting using anti-pdm09 HA sera. The western blot gave a clear readout for HA1 content, confirmed the mobility shift upon de-glycosylation and showed increased amounts of HA1 in the 6:2 FS samples (Figure 6D lower panel). Quantitative measurements of the de-glycosylated samples showed that the 6:2 FS virus gave 1.9-fold greater HA1 yield than the WT reassortant. To test the reproducibility of this finding, HA1 yield was assessed by densitometry of de-glycosylated HA1 following SDS-PAGE and western blot for partially purified virus from 9 independent experiments with the Cal7 and Eng195 reassortants. When examined as paired observations, it was evident that in 8 of the 9 experiments, the FS viruses gave greater HA yields than the parental virus, with only one experiment producing a lower amount (Figure 6E). In one large-scale experiment, the HA1 yield of 6:2 FS was approximately 20-fold higher compared to its 6:2 counterpart. However, in all other experiments, the 6:2 FS virus gave between 1.5 and 3-fold increases in HA1 yield when compared with the 6:2 virus. When the outlier was discounted (as possibly resulting from an artefactually low recovery for the WT sample), average HA1 yield from the other 8 experiments showed 1.9- and 2.4-fold improvements with the segment 3 FS mutation for Cal7 and Eng195 respectively (Table 1).

The HA yield of CVVs with pdm09 glycoproteins has been shown to be improved by engineering chimeric HA genes which contain signal peptide and transmembrane domain/cytoplasmic tail sequences from PR8 HA and the antigenic region of the HA gene from Cal7 (19, 20). To test if these gains were additive with those seen with the FS mutation, we introduced the NIBRG-119 construct, which is a segment 4 with the ectodomain coding region of Cal7 HA and all other sequences (3′- and 5′-noncoding regions, signal peptide, transmembrane domain, and cytoplasmic tail) from PR8 (19) into 6:2 CVV mimics with the WT A(H1N1)pdm09 NA gene and a PR8 backbone with or without the PA-X mutation. Viruses bearing the NIBRG-119 HA did not agglutinate chicken red blood cells (data not shown) so HA yield from eggs was assessed by SDS-PAGE and western blot of partially purified virus. Chimeric HA viruses containing the FS backbone showed an average HA yield improvement of 1.54-fold compared to the WT backbone counterpart, across independent small- and large-scale experiments (Table 1). Thus, the FS mutation is compatible with other rational strategies for increasing egg-grown reverse genetics vaccines.

Following on from this, several pairs of CVV mimics were made with glycoproteins from different IAV strains with either WT or FS mutant PR8 segment 3. These included viruses with glycoproteins of potentially pandemic strains such as highly pathogenic avian virus A/turkey/Turkey/1/2005 (H5N1), as well as low pathogenic avian strains A/mallard/Netherlands/12/2000 (H7N3), A/chicken/Pakistan/UDL-01/2008 (H9N2) and A/mallard/Netherlands/10/99 (H1N1), as well as the human H3N2 strain, A/Hong Kong/1/68, and an early seasonal H3N2 isolate, A/Udorn/307/72 (Table 1). HA yield in eggs was assessed from both the small-scale and large-scale experimental conditions described earlier, by measuring HA titre and HA1 yield from partially purified virus particles. In general, the two techniques were in agreement (Table 1). Ablating PA-X expression moderately improved HA1 yields of some of the CVVs tested: 1.5-fold for the avian H7N3 strain, A/mallard/Netherlands/12/2000 and 1.3-fold for the human H3N2 A/Udorn/307/72 strain. Other CVVs showed lesser or effectively no increases. However, in no case, did ablation of PA-X appear to be detrimental to the growth of CVVs.

## Discussion

Here we show that ablating expression of PA-X resulted in reduced pathogenicity in the chicken embryo model despite the PR8 PA-X protein having relatively low host cell shut-off activity compared to PA-X from other IAV strains. Although loss of PA-X expression had no effect on infectious titres in eggs, subtle differences in virion composition were observed, and more importantly, the HA yield from poor growing 6:2 reassortant vaccine analogues containing the HA and NA segments from A(H1N1) pdm09 strains was significantly improved.

The majority of studies examining the effect of loss of PA-X expression on IAV pathogenicity have used mice as the experimental system. As discussed above, in most cases, the outcome has been increased virulence (30, 34-40), but several studies have found the opposite effect, with PA-X deficiency reducing pathogenicity in mice (37, 41, 42). In adult bird challenge systems using chickens and ducks infected with a highly pathogenic H5N1 virus, abrogating PA-X expression caused increased virulence (35). In our infection model of embryonated hens’ eggs, loss of PA-X expression markedly reduced the pathogenicity in chick embryos. Thus like PB1-F2, another *trans*-frame encoded IAV accessory protein (51), the impact of PA-X expression on viral pathogenicity seems to vary according to both host and virus strain, but not in a fashion that can simply be correlated with mammalian or avian settings.

In previous studies, changes in virulence phenotypes following loss of PA-X expression have been associated with its host cell shut-off function. In the virus strains used, whether from high pathogenicity or low pathogenicity IAV strains, the PA-X polypeptides were shown to significantly affect host cell gene expression. Here, despite PR8 PA-X failing to repress cellular gene expression, a strong phenotypic effect was seen in chicken embryos following loss of PA-X expression. Furthermore, these effects on pathogenicity were more pronounced in an otherwise WT PR8 virus than in a 7:1 reassortant with segment 3 from the highly pathogenic H5N1 avian influenza T/E strain which encodes a PA-X with strong host cell shut-off activity. This lack of correlation between repression of cellular gene expression in avian cells and phenotypic effects in chicken embryos suggests that the PR8 PA-X protein may harbour a function unrelated to host cell shut-off. The PR8 PA-X protein has been proposed to inhibit stress granule formation, but via a mechanism linked to its endonuclease activity and therefore presumably reflecting shut-off activity (52). Alternatively, it could be that the PR8 PA-X polypeptide only exhibits repressive function in specific cell types, such as those of the chorioallantoic membrane (the primary site of virus replication in eggs) or the chick embryo itself. However, since we found low shut-off activity from it in a variety of cells from different species and conversely, no great cell specificity of high activity PA-X polypeptides (data not shown), we do not favour this hypothesis.

Several studies have found that sequences in the X-ORF make positive contributions to the shut-off activity of PA-X (30, 37, 39, 45, 46). In contrast, here we found that for both PR8 and T/E strains of the polypeptide, removal of X-ORF sequences actually increased shut-off activity compared to the WT polypeptide. The effect was relatively modest and in the case of PR8, did not confer equivalent activity to the full-length avian virus PA-X polypeptides (Figure 1C). A similar outcome of greater inhibition from a truncated PA-X polypeptide was seen with a triple reassortant swine influenza virus (42), suggesting that the X-ORF can harbour negative as well as positive regulatory polymorphisms.

In some but not all studies, effects of PA-X mutations on viral pathogenicity have been associated with differences in virus replication *in vivo*. While Jagger et al., (30) did not attribute the increased virulence in mice upon loss of 1918 H1N1 PA-X to virus replication, Gao and colleagues found that increased virulence in mice on loss of H5N1 PA-X was associated with increased titres of ΔPA-X viruses in the lungs, brains and blood of infected mice (34, 39). Similarly, Hu et al. found that increased virulence in chicken, ducks and mice of a ΔPA-X H5N1 virus was associated with increased virus titres in the host (35). Given the postulated role of PA-X-mediated repression of cellular gene expression in controlling host responses to infection, it is reasonable to hypothesise that these differing outcomes reflect the variable interplay between host and virus that is well known to tip in favour of one or other depending on exact circumstance (53). Our present study, where loss of a PA-X with little apparent ability to modulate host gene expression had no significant effect on virus titres in allantoic fluid or the chick embryos themselves, but nevertheless reduced pathogenicity, do not support this hypothesis. However, differences in progeny virion composition in the form of altered ratios of HA to NP and M1 between WT and FS viruses were seen. This may differentially affect their ability to infect specific cell types, as the amount of virus receptor varies between different tissue types and is a known determinant of tissue tropism of influenza viruses (reviewed in (54, 55)).

Our findings have direct implications for HA yield of vaccine viruses in eggs. Ablating PA-X expression did not affect yield from eggs of high growth viruses such as PR8 or 6:2 reassortant CVV mimics containing glycoproteins of human H3N2 strains, or potentially pandemic low pathogenicity avian H9N2 or H1N1 viruses. However, mutation of the PR8 PA-X gene in the background of a CVV analogue containing the HA and NA segments from poor growing strains, such as A(H1N1)pdm09 viruses or a potentially pandemic avian H7N3 isolate, increased HA yield by around 2-fold. The mechanism of improved yield of certain virus subtypes but not others on loss of PA-X expression is unclear. Others have found that mutating the FS site of PR8 PA-X has subtle effects on viral protein expression *in vitro*, including lower levels of M1 (45), perhaps explaining the changes in HA to M1 ratio we see. Beneficial outcomes to HA yield may only be apparent in low-yielding strains where perhaps viral rather than cellular factors are limiting. Alternatively, changes in virion composition between WT and FS viruses could result in subtype/strain-specific effects depending on the balance between HA and NA activities (56). Whatever the mechanism, in no case was loss of PA-X expression detrimental to yield of CVVs, when assessing HA yield of a wide range of different influenza A subtypes/strains. This approach of modifying the PR8 donor backbone therefore potentially supplies a ‘universal’ approach that can be applied to all CVVs that is additive with, but without the need for, generation and validation of, subtype/strain-specific constructs, as is required for strategies based on altering the glycoprotein genes. This could be beneficial to improve antigen yield in a pandemic setting where manufacturers are required to produce large amounts of vaccine quickly.

## Materials and methods

### Cell lines and plasmids

Human embryonic kidney (293T) cells, canine kidney Madin-Darby canine kidney epithelial cells (MDCK) and MDCK-SIAT1 (stably transfected with the cDNA of human 2,6-sialtransferase; (57)) cells were obtained from the Crick Worldwide Influenza Centre, The Francis Crick Institute, London. QT-35 (Japanese quail fibrosarcoma; (58)) cells were obtained from Dr Laurence Tiley, University of Cambridge. Cells were cultured in DMEM (Sigma) containing 10% (v/v) FBS, 100 U/mL penicillin/streptomycin and 100 U/mL GlutaMAX with 1 mg/ml Geneticin as a selection marker for the SIAT cells. Infection was carried out in serum-free DMEM containing 100 U/mL penicillin/streptomycin, 100 U/mL GlutaMAX and 0.14% (w/v) BSA. Cells were incubated at 37°C, 5% CO_2_. Reverse genetics plasmids were kindly provided by Professor Ron Fouchier (A/Puerto Rico/8/34; (59)), Professor Wendy Barclay (A/England/195/2009 (60) and A/turkey/England/50-92/91 (61)), Dr John McCauley (A/California/07/2009; (62)), Dr Laurence Tiley (A/mallard/Netherlands/10/1999 (63)), Professor Robert Lamb (A/Udorn/307/72 (64)), Professor Earl Brown (A/Hong Kong/1/68 (65)) and Professor Munir Iqbal (A/chicken/Pakistan/UDL-01/2008 (66)). RG plasmids for A/mallard/Netherlands/12/2000 (NIBRG-60) and A/turkey/Turkey/1/2005 (NIBRG-23; with the multi-basic cleavage site removed (67)) were made by amplifying HA and NA genes by PCR from cDNA clones available within NIBSC and cloning into pHW2000 vector using BsmB1 restriction sites. A plasmid containing the *Renilla* luciferase gene behind the simian virus 40 early promoter (pRL) was supplied by Promega Ltd.

### Antibodies and sera

Primary antibodies used were: rabbit polyclonal antibody anti-HA for swine H1 (Ab91641, AbCam), rabbit polyclonal anti-HA for H7N7 A/chicken/MD/MINHMA/2004 (IT-003-008, Immune Tech Ltd), mouse monoclonal anti-HA for H5N1 (8D2, Ab82455, AbCam), laboratory-made rabbit polyclonal anti-NP (2915) (68), anti-PA residues 16-213 (expressed as a fusion protein with ß-galactosidase (69), anti-puromycin mouse monoclonal antibody (Millipore MABE343), rabbit anti-PR8 PA-X peptide (residues 211-225) antibody (30) and anti-tubulin-α rat monoclonal antibody (Serotec MCA77G). Secondary antibodies used were: for immunofluorescence, Alexa fluor donkey anti-rabbit IgG 488 or 594 conjugates (Invitrogen), for immunohistochemistry, goat anti-mouse horseradish peroxidase (Biorad 172-1011) and goat anti-rabbit horseradish peroxidase (Biorad 172-1019), for western blot, donkey anti-rabbit IgG Dylight800 or Alexa fluor 680-conjugated donkey anti-mouse IgG (Licor Biosciences).

### Site–directed mutagenesis

The QuikChange® Lightning site-directed mutagenesis kit (Stratagene) was used according to the manufacturer’s instructions. Primers used for site-directed mutagenesis of the segment 3 gene were designed using the primer design tool from Agilent technologies. The strategies used to disrupt the frameshift site (FS) as well as generating C-terminally truncated versions of PA-X via PTCs were as described (30); the cited study used the PTC1 construct.

### Protein analyses

Coupled *in vitro* transcription–translation reactions were carried out in rabbit reticulocyte lysate supplemented with ^35^S-methionine using the Promega TNT system according to the manufacturer’s instructions. SDS–PAGE followed by autoradiography was performed according to standard procedures. Immunoprecipitations were performed as previously described (70). Transfection-based reporter assays to assess host cell shut-off by PA-X (described previously (30)) were performed by co-transfecting QT-35 cells with a reporter plasmid containing the *Renilla* luciferase gene along with pHW2000 plasmids expressing the appropriate segment 3 genes with or without the desired PA-X mutations. 48 h post-transfection, cells were lysed and luciferase activity measured on a Promega GloMax 96-well Microplate luminometer using the Promega *Renilla* Luciferase system.

### Reverse genetics rescue of viruses

All viruses used in this study were made by reverse genetics. 293T cells were transfected with eight pHW2000 plasmids each encoding one of the influenza segments using Lipofectamine(tm) 2000 (Invitrogen). Cells were incubated for 6 hours post-transfection before medium was replaced with DMEM serum-free virus growth medium. At 2 days post-transfection, 0.5 µg/ml TPCK trypsin (Sigma) was added to cells. Cell culture supernatants were harvested at 3 days post-transfection. 293T cell culture supernatants were clarified and used to infect 10-11 day-old embryonated hens’ eggs. At 3 days p.i., eggs were chilled overnight and virus stocks were partially sequenced to confirm identity.

### RNA extraction, RT-PCR and sequence analysis

Viral RNA extractions were performed using the QIAamp viral RNA mini kit with on-column DNase digestion (QIAGEN). Reverse transcription used the influenza A Uni12 primer (AGCAAAAGCAGG) using a Verso cDNA kit (Thermo Scientific). PCR reactions were performed using Pfu Ultra II fusion 145 HS polymerase (Stratagene) or Taq Polymerase (Invitrogen) according to the manufacturers’ protocols. PCR products were purified for sequencing by Illustra GFX PCR DNA and Gel Band Purification kit (GE Healthcare). Primers and purified DNA were sent to GATC biotech for Sanger sequencing (Lightrun method). Sequences were analysed using the DNAstar software.

### Virus titration

Plaque assays, TCID_50_ assays and haemagglutination assays were performed according to standard methods (71). MDCK or MDCK-SIAT cells were used for infectious virus titration, and infectious foci were visualised by either toluidine blue or immunostaining for influenza NP and visualising using a tetra-methyl benzidine (TMB) substrate.

### Virus purification and analysis

Allantoic fluid was clarified by centrifugation twice at 6,500 x g for 10 mins. Virus was then partially purified by ultracentrifugation at 128,000 x g for 1.5 hours at 4°C through a 30% sucrose cushion. For further purification, virus pellets were resuspended in PBS, loaded onto 15-60% sucrose/PBS density gradients and centrifuged at 210,000 x g for 40 mins at 4°C. Virus bands were extracted from gradients and virus was pelleted by ultracentrifugation at 128,000 x g for 1.5 hours at 4°C. Pellets were resuspended in PBS and aliquots treated with N-glycosidase F (New England Biolabs), according to the manufacturer’s protocol. Virus pellets were lysed in Laemmli sample buffer and separated by SDS-PAGE on 10% or 12% polyacrylamide gels under reducing conditions. Protein bands were visualised by Coomassie blue staining (Imperial™ protein stain, Thermo Scientific) or detected by immunostaining in western blot. Coomassie stained gels were scanned and bands quantified using ImageJ software. Western blots were scanned on a Li-Cor Odyssey Infrared Imaging system v1.2 after staining with the appropriate antibodies and bands were quantified using ImageStudio Lite software (Odyssey).

### Chick embryo pathogenesis model

Ten-day old embryonated hens’ eggs were inoculated via the allantoic cavity route with 1000 PFU in 100 μl per egg or mock (serum-free medium only) infected. Embryo viability was subsequently determined by examination of veins lining the shell (which collapse on death) and embryo movement (for a few minutes). At 2 - 3 days p.i. (depending on experiment), embryos were killed by chilling, washed several times in PBS and then scored blind for overt pathology by two observers in each experiment. Scores were 0 = normal, 1 = intact but with dispersed haemorrhages, 2 = small, fragile embryo with dispersed haemorrhages. For histology, embryos were decapitated, washed several times in PBS, imaged and fixed for several days in 4% formalin in PBS. Two embryos per virus condition were sectioned longitudinally and mounted onto paraffin wax. Tissue sections were cut and mounted onto slides and stained with haematoxylin and eosin (H&E) by the Easter Bush Pathology Service. Further sections were examined by immunohistofluorescence performed for influenza NP (62). Sections were deparaffinised and rehydrated and heat-induced antigen retrieval was performed using sodium citrate buffer (10 mM sodium citrate, 0.05% Tween20, pH 6.0). Sections were stained with anti-NP antibody followed by an Alexa fluor-conjugated secondary antibody. Pre-immune bleed serum was also used to confirm specificity of staining by anti-NP antibody. Sections were mounted using ProLong Gold anti-fade reagent containing DAPI (Invitrogen). Stained tissue sections were scanned using a Nanozoomer XR (Hamamatsu) using brightfield or fluorescence settings. Images were analysed using the NDP view 2.3 software (Hamamatsu).

### Graphs and statistical analyses

All graphs were plotted and statistical analyses (Mantel-Cox test, t-tests and Dunnett’s and Tukey’s tests as part of one-way Anova) performed using Graphpad Prism software.

## Acknowledgements

We thank Dr. Francesco Gubinelli, Dr. Carolyn Nicolson and Dr. Ruth Harvey at the Influenza Resource Centre, National Institute for Biological Standards and Control, U. K for their support during experiments performed in their lab, and staff at the Easter Bush Pathology service for pathology support, Bob Fleming and Dr José Pereira for imaging assistance, and Dr. Liliane Chung and. Dr. Marlynne Quigg-Nicol for technical advice.

## References

1. Johnson NPAS, Mueller J. 2002. Updating the Accounts: Global Mortality of the 1918-1920 “Spanish” Influenza Pandemic. Bulletin of the History of Medicine 76:105–115.

2. Dawood FS, Iuliano AD, Reed C, Meltzer MI, Shay DK, Cheng P-Y, Bandaranayake D, Breiman RF, Brooks WA, Buchy P, Feikin DR, Fowler KB, Gordon A, Hien NT, Horby P, Huang QS, Katz MA, Krishnan A, Lal R, Montgomery JM, Mølbak K, Pebody R, Presanis AM, Razuri H, Steens A, Tinoco YO, Wallinga J, Yu H, Vong S, Bresee J, Widdowson M-A. 2012. Estimated global mortality associated with the first 12 months of 2009 pandemic influenza A H1N1 virus circulation: a modelling study. The Lancet Infectious Diseases 12:687–695.

3. Khaperskyy DA, McCormick C. 2015. Timing Is Everything: Coordinated Control of Host Shutoff by Influenza A Virus NS1 and PA-X Proteins. J Virol 89:6528–6531.

4. Vasin AV, Temkina OA, Egorov VV, Klotchenko SA, Plotnikova MA, Kiselev OI. 2014. Molecular mechanisms enhancing the proteome of influenza A viruses: an overview of recently discovered proteins. Virus Res 185:53–63.

5. Palese P, Schulman JL. 1976. Mapping of the influenza virus genome: identification of the hemagglutinin and the neuraminidase genes. Proc Natl Acad Sci U S A 73:2142–2146.

6. Ritchey MB, Palese P, Schulman JL. 1976. Mapping of the influenza virus genome. III. Identification of genes coding for nucleoprotein, membrane protein, and nonstructural protein. J Virol 20:307–313.

7. Neumann G, Watanabe T, Ito H, Watanabe S, Goto H, Gao P, Hughes M, Perez DR, Donis R, Hoffmann E, Hobom G, Kawaoka Y. 1999. Generation of influenza A viruses entirely from cloned cDNAs. Proc Natl Acad Sci U S A 96:9345–9350.

8. Hoffmann E, Neumann G, Kawaoka Y, Hobom G, Webster RG. 2000. A DNA transfection system for generation of influenza A virus from eight plasmids. Proc Natl Acad Sci U S A 97:6108–6113.

9. Fodor E, Devenish L, Engelhardt OG, Palese P, Brownlee GG, Garcia-Sastre A. 1999. Rescue of influenza A virus from recombinant DNA. J Virol 73:9679–9682.

10. Robertson JS, Nicolson C, Harvey R, Johnson R, Major D, Guilfoyle K, Roseby S, Newman R, Collin R, Wallis C, Engelhardt OG, Wood JM, Le J, Manojkumar R, Pokorny BA, Silverman J, Devis R, Bucher D, Verity E, Agius C, Camuglia S, Ong C, Rockman S, Curtis A, Schoofs P, Zoueva O, Xie H, Li X, Lin Z, Ye Z, Chen LM, O’Neill E, Balish A, Lipatov AS, Guo Z, Isakova I, Davis CT, Rivailler P, Gustin KM, Belser JA, Maines TR, Tumpey TM, Xu X, Katz JM, Klimov A, Cox NJ, Donis RO. 2011. The development of vaccine viruses against pandemic A(H1N1) influenza. Vaccine 29:1836–1843.

11. Johnson A, Chen LM, Winne E, Santana W, Metcalfe MG, Mateu-Petit G, Ridenour C, Hossain MJ, Villanueva J, Zaki SR, Williams TL, Cox NJ, Barr JR, Donis RO. 2015. Identification of Influenza A/PR/8/34 Donor Viruses Imparting High Hemagglutinin Yields to Candidate Vaccine Viruses in Eggs. PloS One 10:e0128982.

12. Ping J, Lopes TJ, Nidom CA, Ghedin E, Macken CA, Fitch A, Imai M, Maher EA, Neumann G, Kawaoka Y. 2015. Development of high-yield influenza A virus vaccine viruses. Nat Commun 6:8148.

13. Barman S, Krylov PS, Turner JC, Franks J, Webster RG, Husain M, Webby RJ. 2017. Manipulation of neuraminidase packaging signals and hemagglutinin residues improves the growth of A/Anhui/1/2013 (H7N9) influenza vaccine virus yield in eggs. Vaccine 35:1424–1430.

14. Adamo JE, Liu T, Schmeisser F, Ye Z. 2009. Optimizing viral protein yield of influenza virus strain A/Vietnam/1203/2004 by modification of the neuraminidase gene. J Virol 83:4023–4029.

15. Pan W, Dong Z, Meng W, Zhang W, Li T, Li C, Zhang B, Chen L. 2012. Improvement of influenza vaccine strain A/Vietnam/1194/2004 (H5N1) growth with the neuraminidase packaging sequence from A/Puerto Rico/8/34. Hum Vaccin Immunother 8:252–259.

16. Jing X, Phy K, Li X, Ye Z. 2012. Increased hemagglutinin content in a reassortant 2009 pandemic H1N1 influenza virus with chimeric neuraminidase containing donor A/Puerto Rico/8/34 virus transmembrane and stalk domains. Vaccine 30:4144–4152.

17. Harvey R, Nicolson C, Johnson RE, Guilfoyle KA, Major DL, Robertson JS, Engelhardt OG. 2010. Improved haemagglutinin antigen content in H5N1 candidate vaccine viruses with chimeric haemagglutinin molecules. Vaccine 28:8008–8014.

18. Harvey R, Johnson RE, MacLellan-Gibson K, Robertson JS, Engelhardt OG. 2014. A promoter mutation in the haemagglutinin segment of influenza A virus generates an effective candidate live attenuated vaccine. Influenza Other Respir Viruses 8:605–612.

19. Harvey R, Guilfoyle KA, Roseby S, Robertson JS, Engelhardt OG. 2011. Improved antigen yield in pandemic H1N1 (2009) candidate vaccine viruses with chimeric hemagglutinin molecules. J Virol 85:6086–6090.

20. Medina J, Boukhebza H, De Saint Jean A, Sodoyer R, Legastelois I, Moste C. 2015. Optimization of influenza A vaccine virus by reverse genetic using chimeric HA and NA genes with an extended PR8 backbone. Vaccine 33:4221–4227.

21. Plant EP, Ye Z. 2015. Chimeric neuraminidase and mutant PB1 gene constellation improves growth and yield of H5N1 vaccine candidate virus. J Gen Virol 96:752–755.

22. Plant EP, Liu TM, Xie H, Ye Z. 2012. Mutations to A/Puerto Rico/8/34 PB1 gene improves seasonal reassortant influenza A virus growth kinetics. Vaccine 31:207–212.

23. Cobbin JC, Verity EE, Gilbertson BP, Rockman SP, Brown LE. 2013. The source of the PB1 gene in influenza vaccine reassortants selectively alters the hemagglutinin content of the resulting seed virus. J Virol 87:5577–5585.

24. Cobbin JC, Ong C, Verity E, Gilbertson BP, Rockman SP, Brown LE. 2014. Influenza virus PB1 and neuraminidase gene segments can cosegregate during vaccine reassortment driven by interactions in the PB1 coding region. J Virol 88:8971–8980.

25. Wanitchang A, Kramyu J, Jongkaewwattana A. 2010. Enhancement of reverse genetics-derived swine-origin H1N1 influenza virus seed vaccine growth by inclusion of indigenous polymerase PB1 protein. Virus Res 147:145–148.

26. Gomila RC, Suphaphiphat P, Judge C, Spencer T, Ferrari A, Wen Y, Palladino G, Dormitzer PR, Mason PW. 2013. Improving influenza virus backbones by including terminal regions of MDCK-adapted strains on hemagglutinin and neuraminidase gene segments. Vaccine 31:4736–4743.

27. Giria M, Santos L, Louro J, Rebelo de Andrade H. 2016. Reverse genetics vaccine seeds for influenza: Proof of concept in the source of PB1 as a determinant factor in virus growth and antigen yield. Virology 496:21–27.

28. Mostafa A, Kanrai P, Ziebuhr J, Pleschka S. 2016. The PB1 segment of an influenza A virus H1N1 2009pdm isolate enhances the replication efficiency of specific influenza vaccine strains in cell culture and embryonated eggs. J Gen Virol 97:620–631.

29. Gilbertson B, Zheng T, Gerber M, Printz-Schweigert A, Ong C, Marquet R, Isel C, Rockman S, Brown L. 2016. Influenza NA and PB1 Gene Segments Interact during the Formation of Viral Progeny: Localization of the Binding Region within the PB1 Gene. Viruses 8:238.

30. Jagger BW, Wise HM, Kash JC, Walters KA, Wills NM, Xiao YL, Dunfee RL, Schwartzman LM, Ozinsky A, Bell GL, Dalton RM, Lo A, Efstathiou S, Atkins JF, Firth AE, Taubenberger JK, Digard P. 2012. An overlapping protein-coding region in influenza A virus segment 3 modulates the host response. Science 337:199– 204.

31. Shi M, Jagger BW, Wise HM, Digard P, Holmes EC, Taubenberger JK. 2012. Evolutionary conservation of the PA-X open reading frame in segment 3 of influenza A virus. J Virol 86:12411–12413.

32. Yewdell JW, Ince WL. 2012. Virology. Frameshifting to PA-X influenza. Science 337:164–165.

33. Desmet EA, Bussey KA, Stone R, Takimoto T. 2013. Identification of the N-terminal domain of the influenza virus PA responsible for the suppression of host protein synthesis. J Virol 87:3108–3118.

34. Gao H, Sun Y, Hu J, Qi L, Wang J, Xiong X, Wang Y, He Q, Lin Y, Kong W, Seng LG, Sun H, Pu J, Chang KC, Liu X, Liu J. 2015. The contribution of PA-X to the virulence of pandemic 2009 H1N1 and highly pathogenic H5N1 avian influenza viruses. Sci Rep 5:8262.

35. Hu J, Mo Y, Wang X, Gu M, Hu Z, Zhong L, Wu Q, Hao X, Hu S, Liu W, Liu H, Liu X, Liu X. 2015. PA-X decreases the pathogenicity of highly pathogenic H5N1 influenza A virus in avian species by inhibiting virus replication and host response. J Virol 89:4126–4142.

36. Hayashi T, MacDonald LA, Takimoto T. 2015. Influenza A Virus Protein PA-X Contributes to Viral Growth and Suppression of the Host Antiviral and Immune Responses. J Virol 89:6442–6452.

37. Lee J, Yu H, Li Y, Ma J, Lang Y, Duff M, Henningson J, Liu Q, Li Y, Nagy A, Bawa B, Li Z, Tong G, Richt JA, Ma W. 2017. Impacts of different expressions of PA-X protein on 2009 pandemic H1N1 virus replication, pathogenicity and host immune responses. Virology 504:25–35.

38. Hu J, Mo Y, Gao Z, Wang X, Gu M, Liang Y, Cheng X, Hu S, Liu W, Liu H, Chen S, Liu X, Peng D, Liu X. 2016. PA-X-associated early alleviation of the acute lung injury contributes to the attenuation of a highly pathogenic H5N1 avian influenza virus in mice. Med Microbiol Immunol 205:381–395.

39. Gao H, Sun H, Hu J, Qi L, Wang J, Xiong X, Wang Y, He Q, Lin Y, Kong W, Seng LG, Pu J, Chang KC, Liu X, Liu J, Sun Y. 2015. Twenty amino acids at the C-terminus of PA-X are associated with increased influenza A virus replication and pathogenicity. J Gen Virol 96:2036–2049.

40. Gao H, Xu G, Sun Y, Qi L, Wang J, Kong W, Sun H, Pu J, Chang KC, Liu J. 2015. PA-X is a virulence factor in avian H9N2 influenza virus. J Gen Virol 96:2587– 2594.

41. Nogales A, Rodriguez L, DeDiego ML, Topham DJ, Martinez-Sobrido L. 2017. Interplay of PA-X and NS1 Proteins in Replication and Pathogenesis of a Temperature-Sensitive 2009 Pandemic H1N1 Influenza A Virus. J Virol 91:e00720–00717.

42. Xu G, Zhang X, Liu Q, Bing G, Hu Z, Sun H, Xiong X, Jiang M, He Q, Wang Y, Pu J, Guo X, Yang H, Liu J, Sun Y. 2017. PA-X protein contributes to virulence of triple-reassortant H1N2 influenza virus by suppressing early immune responses in swine. Virology 508:45–53.

43. Naffakh N, Massin P, van der Werf S. 2001. The transcription/replication activity of the polymerase of influenza A viruses is not correlated with the level of proteolysis induced by the PA subunit. Virology 285:244–252.

44. Hayashi T, Chaimayo C, McGuinness J, Takimoto T, Abdel-Wahab M. 2016. Critical Role of the PA-X C-Terminal Domain of Influenza A Virus in Its Subcellular Localization and Shutoff Activity. Journal of Virology 90:7131–7141.

45. Khaperskyy DA, Schmaling S, Larkins-Ford J, McCormick C, Gaglia MM. 2016. Selective Degradation of Host RNA Polymerase II Transcripts by Influenza A Virus PA-X Host Shutoff Protein. PloS Pathog 12:e1005427.

46. Oishi K, Yamayoshi S, Kawaoka Y. 2015. Mapping of a Region of the PA-X Protein of Influenza A Virus That Is Important for Its Shutoff Activity. J Virol 89:8661–8665.

47. Firth AE, Jagger BW, Wise HM, Nelson CC, Parsawar K, Wills NM, Napthine S, Taubenberger JK, Digard P, Atkins JF. 2012. Ribosomal frameshifting used in influenza A virus expression occurs within the sequence UCC_UUU_CGU and is in the +1 direction. Open Biol 2:120109.

48. Abernathy E, Clyde K, Yeasmin R, Krug LT, Burlingame A, Coscoy L, Glaunsinger B. 2014. Gammaherpesviral gene expression and virion composition are broadly controlled by accelerated mRNA degradation. PloS Pathog 10:e1003882.

49. Abt M, de Jonge J, Laue M, Wolff T. 2011. Improvement of H5N1 influenza vaccine viruses: influence of internal gene segments of avian and human origin on production and hemagglutinin content. Vaccine 29:5153–5162.

50. Harvey R, Wheeler JX, Wallis CL, Robertson JS, Engelhardt OG. 2008. Quantitation of haemagglutinin in H5N1 influenza viruses reveals low haemagglutinin content of vaccine virus NIBRG-14 (H5N1). Vaccine 26:6550–6554.

51. Kamal RP, Alymova IV, York IA. 2017. Evolution and Virulence of Influenza A Virus Protein PB1-F2. Int J Mol Sci 19:96.

52. Khaperskyy DA, Emara MM, Johnston BP, Anderson P, Hatchette TF, McCormick C. 2014. Influenza a virus host shutoff disables antiviral stress-induced translation arrest. PloS Pathog 10:e1004217.

53. Newton AH, Cardani A, Braciale TJ. 2016. The host immune response in respiratory virus infection: balancing virus clearance and immunopathology. Semin Immunopathol 38:471–482.

54. Klenk HD, Garten W, Matrosovich M. 2011. Molecular mechanisms of interspecies transmission and pathogenicity of influenza viruses: Lessons from the 2009 pandemic. Bioessays 33:180–188.

55. Baigent SJ, McCauley JW. 2003. Influenza type A in humans, mammals and birds: determinants of virus virulence, host-range and interspecies transmission. Bioessays 25:657–671.

56. Benton DJ, Martin SR, Wharton SA, McCauley JW. 2015. Biophysical measurement of the balance of influenza a hemagglutinin and neuraminidase activities. J Biol Chem 290:6516–6521.

57. Matrosovich M, Matrosovich T, Carr J, Roberts NA, Klenk HD. 2003. Overexpression of the alpha-2,6-sialyltransferase in MDCK cells increases influenza virus sensitivity to neuraminidase inhibitors. J Virol 77:8418–8425.

58. Moscovici C, Moscovici MG, Jimenez H, Lai MM, Hayman MJ, Vogt PK. 1977. Continuous tissue culture cell lines derived from chemically induced tumors of Japanese quail. Cell 11:95–103.

59. de Wit E, Spronken MI, Bestebroer TM, Rimmelzwaan GF, Osterhaus AD, Fouchier RA. 2004. Efficient generation and growth of influenza virus A/PR/8/34 from eight cDNA fragments. Virus Res 103:155–161.

60. Brookes DW, Miah S, Lackenby A, Hartgroves L, Barclay WS. 2011. Pandemic H1N1 2009 influenza virus with the H275Y oseltamivir resistance neuraminidase mutation shows a small compromise in enzyme activity and viral fitness. J Antimicrob Chemother 66:466–470.

61. Howard W, Hayman A, Lackenby A, Whiteley A, Londt B, Banks J, McCauley J, Barclay W. 2007. Development of a reverse genetics system enabling the rescue of recombinant avian influenza virus A/Turkey/England/50-92/91 (H5N1). Avian Dis 51:393–395.

62. Turnbull ML, Wise HM, Nicol MQ, Smith N, Dunfee RL, Beard PM, Jagger BW, Ligertwood Y, Hardisty GR, Xiao H, Benton DJ, Coburn AM, Paulo JA, Gygi SP, McCauley JW, Taubenberger JK, Lycett SJ, Weekes MP, Dutia BM, Digard P. 2016. Role of the B Allele of Influenza A Virus Segment 8 in Setting Mammalian Host Range and Pathogenicity. J Virol 90:9263–9284.

63. Bourret V, Croville G, Mariette J, Klopp C, Bouchez O, Tiley L, Guérin J-L. 2013. Whole-genome, deep pyrosequencing analysis of a duck influenza A virus evolution in swine cells. Infection, Genetics and Evolution 18:31–41.

64. Chen BJ, Leser GP, Jackson D, Lamb RA. 2008. The influenza virus M2 protein cytoplasmic tail interacts with the M1 protein and influences virus assembly at the site of virus budding. J Virol 82:10059–10070.

65. Ping J, Dankar SK, Forbes NE, Keleta L, Zhou Y, Tyler S, Brown EG. 2010. PB2 and Hemagglutinin Mutations Are Major Determinants of Host Range and Virulence in Mouse-Adapted Influenza A Virus. Journal of Virology 84:10606–10618.

66. Long JS, Giotis ES, Moncorge O, Frise R, Mistry B, James J, Morisson M, Iqbal M, Vignal A, Skinner MA, Barclay WS. 2016. Species difference in ANP32A underlies influenza A virus polymerase host restriction. Nature 529:101–104.

67. Robertson JS, Engelhardt OG. 2010. Developing vaccines to combat pandemic influenza. Viruses 2:532–546.

68. Noton SL, Medcalf E, Fisher D, Mullin AE, Elton D, Digard P. 2007. Identification of the domains of the influenza A virus M1 matrix protein required for NP binding, oligomerization and incorporation into virions. J Gen Virol 88:2280–2290.

69. Blok V, Cianci C, Tibbles KW, Inglis SC, Krystal M, Digard P. 1996. Inhibition of the influenza virus RNA-dependent RNA polymerase by antisera directed against the carboxy-terminal region of the PB2 subunit. J Gen Virol 77:1025–1033.

70. Poole E, Elton D, Medcalf L, Digard P. 2004. Functional domains of the influenza A virus PB2 protein: identification of NP- and PB1-binding sites. Virology 321:120–133.

71. Klimov A, Balish A, Veguilla V, Sun H, Schiffer J, Lu X, Katz JM, Hancock K. 2012. Influenza virus titration, antigenic characterization, and serological methods for antibody detection. Methods Mol Biol 865:25–51.

